# Genotype, nitrogen and herbivory shape plant defense: the case of a vitamin-enriched maize

**DOI:** 10.1101/160333

**Authors:** Agnès Ardanuy, Victoria Pastor, Gaëtan Glauser, Paul Christou, Ted C. J. Turlings, Ramon Albajes

## Abstract

The cultivation of crops with novel traits could interfere with ecosystem services delivered by arthropods through bottom-up effects. Here we tested the hypothesis that a vitamin enriched maize (Carolight^R^) is similar in terms of plant-arthropod interactions to its wild type when compared in controlled environment and under field conditions. In order to assess the robustness of their equivalence we tested two nitrogen availability regimes. We used arthropod field abundance, the behavior and fitness of a keystone maize herbivore - the leafhopper *Zyginidia scutellaris* - and above ground chemistry of maize plants (volatile, hormone and metabolite profiling) as indicators of potential changes in plant-insect interactions. Nitrogen availability was the key driver of herbivore abundance and behavior, and determined direct and indirect chemical defense in maize plants. Both genotypes presented similar constitutive and inducible phytohormone profiles independently of the nitrogen regime. However, feeding by the herbivore suppressed the levels of JA-Ile and JA, without impairing the release of induced plant volatiles. Carolight^R^ and M37W differed to some degree in the concentrations of phenolics (hydroxycinnamic acids and lignans) and in the abundance of a volatile compound. Overall the effect of maize genotype on the herbivores was smaller than the effect of nitrogen fertilization.

**HIGHLIGHT:** We show the separate and interactive effects of nitrogen availability and genotype on the performance and behavior of a herbivore, and related these changes to constitutive and inducible maize defenses.

## INTRODUCTION

One of the issues regarding the cultivation of novel crops (genetically modified or otherwise) is their possible effect on insect biodiversity and associated ecosystem services in agriculture. The mandatory environmental risk assessment for cultivation of novel crops addresses the hypothesis that the traits introduced into the novel crops do not adversely affect the non-target arthropods. Risk assessments are comparative in the sense that novel crops are screened for phenotypic and compositional equivalence to their wild type counterpart, and the biologically meaningful differences observed between them are a consequence of the novel trait (Wolt *et al.*, 2010) and are subsequently evaluated. In addition to risk assessment purposes, studying the herbivore community responses to novel varieties, and in turn the biochemical responses of those novel plant varieties to insect herbivory and nutrient availability may help improve our understanding of plant chemical profiles and their role in plant–herbivore interactions.

An elite South African maize inbred was engineered to deliver pro-vitamin A (and other nutritionally important carotenoids such as lutein, zeaxanthin and lycopene) in the diet and thus address vitamin A and other nutritional deficiencies in at-risk populations in developing countries. The kernels of this novel maize (Carolight^R^) accumulate higher levels of 3 vitamins in the endosperm through the simultaneous engineering of 3 separate metabolic pathways: 169-fold the normal amount of beta-carotene (provitamin A), 6-fold the normal amount of ascorbate (vitamin C), and double the normal amount of folate (vitamin B9) (Naqvi et al., 2009). Molecular and biochemical characterization of Carolight^R^ seeds (transcript, proteome, and metabolite profiles) indicated changes in sugar and lipid metabolism in the endosperm with respective to the wild type due to the higher up-stream metabolite demand by the extended biosynthesis capacities for terpenoids and fatty acids (Decourcelle et al., 2015). Nevertheless under field conditions the metabolic phenotype of vitamin-enriched maize kernels under contrasting soil nitrogen conditions was indistinguishable from the wild type in terms of carotenoid accumulation in leaves, photosynthetic activity, sensitivity to source limitation and yield (Zanga *et al.*, 2016). Authors concluded that the additional metabolic requirements of Carolight^R^ endosperm did not affect agronomic performance. Interestingly gravid females of the key Mediterranean maize pest *Sesamia nonagrioides* preferred the volatiles of the wild type to Carolight^R^ in an olfactometer setting (Cruz and Eizaguirre, 2015), which led to the notion that vitamin enriched maize might modify the outcome of plant-insect interactions.

The strong influence of plant chemical traits on food webs has been demonstrated experimentally both above and below ground (e.g. van der Putten et al. 2001, Ode 2006). As it is not possible to measure all ecological interactions between a plant and its associated insect species, we used the arthropod field abundance, the behaviour and fitness of an herbivore keystone species (Albajes *et al.*, 2011) and above ground chemistry of maize plants as indicators of possible modifications in plant–insect interactions. We therefore tested the hypothesis that Carolight^R^ is similar in terms of plant-arthropod interactions to its wild type line (M37W) when compared in a controlled environment and in the field. The over-arching objectives of the current study were to: (i) determine if Carolight^R^ and M37W influence abundance and dynamics of herbivores and natural enemies in the field; (ii) determine potential impact of both genotypes on herbivore choice and performance under controlled conditions; (iii) characterize the chemical profiles of leaves usually consumed by most herbivores (and thus involved in plant–insect interactions) in both genotypes. Characterization was carried out through volatile, hormone and metabolite profiling. In order to broaden the range of environments in the study and to test the consistency of performance between Carolight^R^ and M37W, we compared both genotypes under different substrate nitrogen availability regimes. The data and conclusions from our studies not only validate the use of plant-insect interactions in the environmental risk assessment of crops with novel traits, but importantly also shed light into the biochemical and metabolic components that underpin the mechanisms involved in maize-insect interactions.

## MATERIAL AND METHODS

### Plants and nitrogen treatments

Seeds of the elite South African maize (Zea mays L.) inbred cv. M37W and its vitamin enriched derived line Carolight^R^ were obtained from the Applied Plant Biotechnology Group at Universitat de Lleida-Agrotecnio Center.

A field experiment was carried out in order to evaluate the performance of Carolight^R^ and M37W in terms of arthropod community composition and dynamics. The experimental design encompassed a factorial combination of the two maize genotypes and two nitrogen treatments. Plots were randomized with four replicates per genotype-nitrogen combination, each consisting of 6 rows, 70 cm apart and 6.47 m in length (approximately 4 plants per meter). Maize was planted on 5 May 2013. Two different fertilization regimes were applied on 9 July 2013: Control = 0 kg ha^-1^ and +N = 200 kg ha^-1^ as urea at the V6 stage (six fully expanded leaves). Each plot was fully irrigated.

For laboratory experiments, seeds from each line were sown in plastic pots (10 cm high, 5 cm diameter) in vermiculite, and germinated in the greenhouse. Forty maize plants (seven to ten days old) were placed in plastic containers and provided 2.5 l of hydroponic solution for 10-12 days. Two hydroponic solutions were tested: a control solution and a solution with an increased content in nitrogen (+N). The control solution consisted of a half-strength modified Hoagland solution with micro-nutrients provided at full strength. The solution with nitrogen (+N) consisted of a control solution in which 8 mM of NH_4_NO_3_ was added. The hydroponic solutions were adjusted to pH 5.9, and were buffered with MES tampon. The solution was replaced every 3-4 days.

### Insects and herbivory treatments

A colony of the leafhopper *Z. scutellaris* was established from small grain cereal and maize fields at the Universitat de Lleida (Spain). The colony was reared under controlled conditions (16:8 h L:D, 24±5 °C) on maize plants (var. Delprim).

Plants were transferred to an experimental chamber equipped with full spectrum light benches (24±2 °C, 40±10% r.h., 16:8 h L/ D, and 8000 lm m^-2^) the day prior the experiments started. Plants used for volatile collection were enclosed in custom made Nalophan bags (Omya AG, Oftringen, Switzerland, 150 mm diameter) closed with a parafilm seal at the top of the plastic pot. Plants used for hormone and non-targeted metabolome profiling were enclosed in bottom cut PET plastic bottles covered with muslin cloth. Herbivore treatment was initiated on the following day by exposing plants to ten *Z. scutellaris* adults for 24 h in the case of volatile analysis and non-targeted metabolome profiling, and 24, 48 and 96h for hormone profiling. The timing was chosen based on a previous study which indicated a strong induction of plant volatiles at 24h after the start of leafhopper feeding (Ardanuy *et al.*, 2016).

### Field herbivore and natural enemy abundance

Visual sampling of arthropod fauna was conducted on whole plants from the 9th of July (V6-7 stage) to the 16th of September, 2013 every other week (5 samplings in total) according to Albajes et al. (2011). We sampled four plants from each plot randomly, and we recorded the number of herbivores and their natural enemies per plant. Herbivore counts were grouped in five taxonomic units: Thysanoptera (thrips), Hemiptera\Aphididae (aphids), Hemiptera\Cicadellidae (leafhoppers, mainly *Zyginidia scutellaris*), and Hemiptera\ Delphacidae (planthoppers, *Laodelphax striatellus*) and Lepidoptera (*Spodoptera spp., Helicoverpa armigera*, corn borers). Later we transformed aphid counts into an abundance scale (0, no aphids; 1, isolated aphids; 2, small colony; 3, medium colony; 4, large colony). Natural enemy counts were grouped in Hemiptera\Anthocoridae, Hemiptera\Miridae, Neuroptera, Coccinellidae, Thysanoptera (thrips) and Arachnida.

We calculated the sum of abundances per plot and sampling date for all taxonomic units. We tested the effects of genotype, nitrogen, and sampling date on herbivore and natural enemy community with a permutational MANOVA using the Adonis function in the package vegan in R (Oksanen *et al.*, 2013). We then performed univariate analysis at the species level for herbivore abundance data with a generalized linear model following a Negative Binomial distribution in which sampling date, nitrogen treatment and genotype and their interactions were used as fixed factors. Aphid abundance was analyzed with an ordinal logistic regression. All statistical analyses were performed using R (R Development Core Team) unless otherwise indicated.

### Herbivore performance and plant choice

Leafhopper performance was tested by transferring 1-day old leafhopper nymphs from the colony to maize plants and letting them develop until adult stage. Plant treatments consisted of a factorial combination of the two maize genotypes and two N treatments (control and +N) (n=13-15 plants per treatment). Plants were enclosed in plastic bottles with their bottom open, covered by cloth to prevent leafhoppers from escaping; each plant contained 3 leafhoppers. Plants were monitored daily until leafhoppers reached adult stage. Leafhoppers were then removed and placed in 0.5 mm eppendorfs and frozen at −20°C until sexed and weighed. When there was more than one leafhopper per sex in a plant we averaged final weight and developmental time. Final weight of leafhopper individuals and developmental time was analyzed with a GLM following a Gaussian distribution using the variables insect sex, nitrogen regime and genotype and their interactions as factors.

The effects of plant volatiles emitted by the different combination of varieties and nitrogen treatments on the behavior of the leafhopper were investigated in a six-arm-olfactometer (for details see Turlings, Davison, & Tamò, 2004). A plant from each genotype-nitrogen treatment was placed in glass vessels one hour before the assay began. Two empty vessels were used as blanks. Purified and humidified air entered each odor source bottle at 0.8 l/min via Teflon tubing (adjusted by a manifold with four flow-meters; Analytical Research System, Gainesville, FL, USA) and carried the volatiles through to the olfactometer compartment. The position of the odor sources in the olfactometer was randomly assigned each experimental day to avoid position-bias.

At least half an hour before the experiment started groups of six *Z. scutellaris* females were isolated in pipette tips by means of a manual aspirator, and covered in parafilm. Twelve leafhoppers were freed at the base of the olfactometer and left for 45 minutes. Only when an insect entered an arm and passed the screw cap fitting or was recovered in the bulb we considered it had made a choice. Three times twelve females were tested per experiment per day. All olfactometer tests were conducted between 10 am and 4 pm under light benches (24±2 °C). Each experiment was performed 7 times on different days. This resulted in 7 independent replicates for each olfactometer setup.

Olfactometer choice counts were analyzed with a GLM following a Poisson distribution, with nitrogen regime and genotype and their interactions as factors. Pair-wise comparisons were performed with using Tukey's HSD.

### Analysis of volatile profiles

VOCs were collected simultaneously from herbivore-damaged plants and from control non-damaged plants for all the treatments consisting of the factorial combination of genotype and nitrogen treatments. Two tubular glass outlets (23x17x12 mm) with a screw cap were attached to the bottom and top of the bag respectively (as described by Turlings et al. 1998). Clean air was supplied to the system through the top outlet via Tygon tubing connected to a flowmeter (Analytical Research Systems) and through the bottom device air was pulled through a volatile adsorbent trap at a rate of 1 l/min using a vacuum pump. We collected volatiles of each odor source for 5h using adsorbent traps consisting of a glass tube (4 mm ID) packed with 25 mg Super-Q polymer (80–100 mesh) (Alltech Associates, Deerfield, Illinois, USA). We performed seven experimental replicates for all treatments on different days.

The traps were then extracted with 150 μl dichloromethane (Suprasolv, Merck, Dietikon, Switzerland), and 200 ng of n-octane and n-nonyl acetate (Sigma, Buchs, Switzerland) in 10 μl dichloromethane were added to the samples as internal standards. Samples were analyzed with a GC-MS as described in Ardanuy et al., (2016). The detected volatiles were identified by comparison of their mass spectra with those of the NIST 05 library and by comparison of retention times with those from a library from earlier assays.

Permutational MANOVA was used to evaluate whether the VOC blend varied between herbivore treatments, nitrogen availability regimes and among genotypes. The abundance of the components of the volatile blend was used as the response variable, while herbivore treatment, nitrogen regime, plant genotype, and their double interactions were used as independent variables. In addition the amount of each individual compound was compared among treatments using a non-parametric Kruskal-Wallis test followed by Dunn’s test.

### Plant hormones and hydroxycinnamic acid analysis

A targeted analysis of plant hormones and phenolic compounds of herbivore-damaged plants (n=3) and control plants (n=3) was performed for each combination of genotype-nitrogen levels at three time points (24, 48 and 96h) after the experiment started. The aboveground part of the plants was flash frozen with liquid nitrogen and stored at −80°C until freeze dried. The experiment was repeated three times. The hormones jasmonoyl-L-isoleucine (JA–Ile), 12-oxo-phytodienoic acid (OPDA), jasmonic acid (JA), salicylic acid (SA), abscisic acid (ABA) and indole-3-acetic acid (IAA), and the hydroxycinnamic acids caffeic acid, chlorogenic acid and ferulic acid were analyzed by ultra-performance liquid chromatography coupled to mass spectrometry (UPLC-MS), as described by Camañes et al. (2012). Data from the three experiments were log-transformed and analyzed by a linear model with nitrogen regime and genotype and their interactions as factors, and experiment as a block. Within an experiment pair-wise comparisons were performed using the Klustal-Wallis test.

### Metabolite fingerprinting

Non-targeted metabolite profiling of herbivore damaged (n=5) and control plants (n=5) was performed for each combination genotype-nitrogen levels. The aboveground part of the plants was flash frozen with liquid nitrogen 24h after the experiment started and stored at −80°C. Each sample was ground to powder using a mortar previously frozen in liquid nitrogen. The frozen powder was weighed (100 mg ±1mg) in an Eppendorf tube, and 500 μl of extraction solvent (MeOH:H_2_O:formic acid 80:20:0.5) and a few glass beads were added. Samples were briefly vortexed and then extracted in a bead mill for three minutes at 30Hz. After centrifugation at 10,000 rpm for 10 min (Hettich mikrolitter D 7200, Buford, GA, USA) the supernatant was transferred to a new Eppendorf tube, to which 350 μl of dichloromethane was added. Samples were vortexed and centrifuged again to separate the two phases. The upper phase was recovered (150 μl) and transferred to HPLC vials.

Metabolite analysis was performed using an Acquity UPLC™ system (Waters) coupled to Synapt G2 QTOF mass spectrometer (Waters) through an electrospray interface (ESI). The separation was performed on an Acquity BEH C18 column (50 × 2.1 mm i.d., 1.7 μm particle size) at a flow rate of 0.6 mL min-1. The injection volume was 3 μl and the autosampler and column temperatures were kept at 15 and 40 °C, respectively. The mobile phase consisted of 0.05% formic acid (FA) in water (phase A) and 0.05% FA in acetonitrile (phase B). The segmented gradient program was as follows: 2% B to 35% B in 3.0 min, 35% B to 100% B in 3.0 min, held at 100% B for 1.5 min, re-equilibrated to initial conditions (2% B) for 1.5 min. Data acquisitions was performed in ESI-negative and ESI-positive modes over a mass range of 100–1000 Da. The MSe mode, in which the instrument alternatively acquires data at low (4 eV; 0.15 s scan time) and high (10-30 eV ramp; 0.15 s scan time) collision energies, was used. The mass spectrometer was internally calibrated by infusing a 500 ng/mL solution of leucine-enkephalin at a flow rate of 15 ul/min through the LockSpray™ probe. The system was controlled by Masslynx v4.1.

Metabolite raw data was transformed to CDF using Databridge provided by the Masslynx package. The CDF data was processed with R for statistical computing using XCMS package for relative quantification (Smith *et al.*, 2006). ESI-negative and ESI-positive data were combined, log-transformed and Pareto scaled prior to analysis. Pareto scaling gives each variable a variance equal to the square root of its standard deviation.

The advantage of using this technique rather than scaling to unit variance is that the former reduces the relative importance of large values but keeps data structure partially intact (van den Berg *et al.*, 2006). First a permutational MANOVA was used to evaluate whether the metabolite fingerprint consistently varied among genotypes, nitrogen availability regimes and herbivore treatments and the influences of the interactions of the factors (permutations=999). Next, a principal component analysis (PCA) was conducted as an unsupervised method to visualize variability and clustering in the data set.

Partial least squares–discriminant analyses (PLS–DA) were performed to identify differently detected ions between plant experimental factors - wild type vs. Carolight^R^, control nitrogen vs. nitrogen treatment, and controls vs. leafhopper-induced plants - given that interactions between factors were non-significant in the perMANOVA. PLS—DA is a supervised multivariate analysis technique, which maximizes the covariance between the X–(spectral intensities) and the Y–matrix (group information). We assessed model reliability using CV-ANOVA. New components were only added to the model when significant according to the cross–validation. R^2^X and R^2^Y represent the fraction of the variance of X and Y matrix, respectively, while Q^2^Y suggests the predictive accuracy of the model. Variable influence on projection (VIP) was used to select the most influential metabolites to group separation in the validated PLS-DA models. The VIP values summarize the overall contribution of each X-variable to the model, summed over all components and weighted according to the Y variation accounted for by each component. The Sum of squares of all VIP's is equal to the number of terms in the model - the average VIP is equal to 1- and thus terms with large VIP are the most important for explaining Y. We considered that metabolites with a VIP> 2 were extremely influential for treatment separation. The ions with VIP>2 for each experimental factor (genotype, nitrogen and herbivory) were screened for putative identification using the pathway tool from MarVis (Kaever *et al.*, 2014). The MS/MS fragmentation of the metabolites was compared with candidate compounds identified in databases or earlier publications, especially when the metabolites were already reported in maize. Metabolite multivariate analysis (PCA, PLS-DA) was performed with SIMCA–P software (v. 11.0, Umetrics, Umeå, Sweden).

## RESULTS

### Effects of genotype and nitrogen on arthropod communities in the field

The most prominent source of variation in insect abundances in the field was plant developmental stage, which reflects seasonal insect dynamics in the plot (Table 1). Thus, abundance of maize herbivores was mainly influenced by the developmental stage of the plant (perMANOVA R^2^ = 0.62, p=<0.001) and to a minor extend by nitrogen regime (R^2^ = 0.13, p=0.085) while no effects were attributable to genotype (R^2^ = 0.07, p=0.213) or genotype x nitrogen interaction (R^2^ = 0.02, p=0.684). Similarly, maize developmental stage was the main factor explaining the variation in the abundance of the natural enemies recorded in the study (R^2^ = 0.28, p=<0.001) whilst genotype and nitrogen were not significant for determining community composition.

**Table 1.** Effects of maize genotype, nitrogen treatment, their interaction and maize’s developmental stage on field abundances of herbivores and natural enemies. Arthropod abundance was determined by visual sampling on five maize developmental stages. Significant effects (α=0.05) appear in bold.

| **Herbivores** |  | **Leafhoppers** |  | **Planthoppers** |  | **Thrips** |  | **Lepidoptera** |  | **Aphids** |  |
| --- | --- | --- | --- | --- | --- | --- | --- | --- | --- | --- | --- |
| *Factors* | df | $\chi^2$ | p | $\chi^2$ | p | $\chi^2$ | p | $\chi^2$ | p | $\chi^2$ | p |
| genotype | 1 | 4.09 | **0.046** | 0.09 | 0.779 | 1.41 | 0.233 | 2.19 | 0.076 | 0.64 | 0.423 |
| nitrogen | 1 | 5.28 | **0.023** | 11.9 | **<0.002** | 0.15 | 0.693 | 0.18 | 0.613 | 10.91 | **0.001** |
| genotype × nitrogen | 1 | 2.1 | 0.150 | 0.43 | 0.547 | 0.45 | 0.501 | 1.38 | 0.161 | 2.57 | 0.109 |
| develop. stage | 4 | 223.14 | **<0.001** | 32.98 | **<0.001** | 157.96 | **<0.001** | 181.53 | **<0.001** | 45.65 | **<0.001** |

| **Natural enemies** |  | **Anthocoridae** |  | **Chrysopidae** |  | **Thrips** |  | **Coccinellidae** |  | **Miridae** |  | **Arachnida** |  |
| --- | --- | --- | --- | --- | --- | --- | --- | --- | --- | --- | --- | --- | --- |
| *Factors* | df | $\chi^2$ | p | $\chi^2$ | p | $\chi^2$ | p | $\chi^2$ | p | $\chi^2$ | p | $\chi^2$ | p |
| genotype | 1 | 0.012 | 0.914 | 0.29 | 0.564 | 0.161 | 0.686 | 0.05 | 0.83 | 1.17 | 0.286 | 1.14 | 0.306 |
| nitrogen | 1 | 0.49 | 0.49 | 0.29 | 0.564 | 0.875 | 0.346 | 0.21 | 0.67 | 0.16 | 0.69 | 1.14 | 0.306 |
| genotype × nitrogen | 1 | 1.95 | 0.168 | 0.02 | 0.881 | 0.24 | 0.619 | 0.15 | 0.72 | 1.90 | 0.175 | 0.00 | 0.992 |
| develop. stage | 4 | 52.41 | **<0.001** | 16.45 | **<0.001** | 45.08 | **<0.001** | 22.68 | **<0.001** | 57.08 | **<0.001** | 16.93 | **0.004** |

Leafhoppers and thrips were the most abundant herbivore taxa in the field, and Anthocoridae and spiders the most abundant natural enemy taxa (Supplementary material, Fig. S.2, S.3, S.4). Univariate analysis revealed that Hemipteran herbivores (leafhoppers, planthoppers and aphids) were more abundant in the higher nitrogen treatments independently of population dynamics (Table 1). Only leafhopper populations were influenced by plant genotype: Carolight^R^ plots supported lower populations of leafhopper nymphs than the wild type (Table 1). Levels of other herbivores such as thrips and Lepidoptera were not influenced by nitrogen treatment or genotype (Table 1). Overall the variation of natural enemy taxa was attributable to population dynamics, and no differences were detected between any of the treatments (Table 1).

### Effects of plant variety and nitrogen levels on herbivore choice and performance

Plants from both genotypes in the high nitrogen hydroponic treatments (+N) were taller and shoots were more robust than plants grown under control nitrogen conditions (Supplementary material Fig. S.1). Genotype and nitrogen factors did not impact herbivore performance as sex was the only significant predictor of final weight (F_1,62_=121.40, p<0.001) and developmental time (F_1,62_=8.71, p=0.032). Overall, plants from both genotypes grown under high nitrogen attracted more female leafhoppers than plants grown with no additional nitrogen (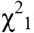= 25.22, p<0.001); however, when considering only the high nitrogen treatment Carolight^R^ was preferred (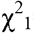= 4.19, p=0.04) (Fig. 2). Leafhoppers chose maize plants over empty bottle control treatments (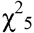= 30.70, p<0.001) a result that validates the experimental setup.

### Effects of plant variety, nitrogen levels and herbivory on volatile compounds

Seven volatile compounds were quantified in our study (Table 2) and all seven had been previously reported in maize (Degen *et al.*, 2004). We expected a small number and amount of volatile compounds in control and herbivore induced plants given that (i) the wild type line M37W produces low amounts of volatile inducible terpenes (Richter *et al.*, 2016) and that (ii) *Z. scutellaris* induced plants do not emit the green leaf volatiles (Z)-3-hexenal and (E)-2-hexenal (Ardanuy *et al.*, 2016). Herbivory explained the most variability in volatile blends (perMANOVA R^2^ =0.647, p=0.001), and a clear separation between control and herbivore induced plants was observed in PC1 (Supplementary material, Fig. S.5). Herbivore damaged plants emitted DMNT, indole, E-β-farnesene and (E)-β-bergamotene in addition to α-copaene, E-β-caryophyllene and β-sesquiphellandrene. However, a significant genotype per nitrogen interaction was detected (perMANOVA R^2^=0.036, p=0.007). In particular, individual differences in volatile emission between nitrogen regimes could be attributed for α-copaene and E-β-caryophyllene (Table 2), while differences between genotypes were only detected for β-sesquiphellandrene in the high nitrogen treatment consistent with the preference of *Z. scutellaris* females for Carolight^R^ +N in the olfactometer assay. An effect of the experimental day of volatile collection was detected on the volatile blend (R^2^=0.028, p=0.011).

**Table 2.** Volatile emissions (ng/h) from control and herbivore-induced maize plants (Carolight^R^, Wild type) at two different N availability treatments (control, +N) (n=7). Amounts of each compound were compared among treatments using a non-parametric Kruskal-Wallis test followed by Dunn’s test (*p<0.05, **p<0.01, ***p<0.001). Compounds denoted with “N” were only tentatively identified by comparison of their MS to that reported in libraries.

| | Wild type | | Carolight | | Wild type+N | | Carolight+N | | $\chi^2$ | P |
| --- | --- | --- | --- | --- | --- | --- | --- | --- | --- | --- |
|  | mean | ±SE | mean | ±SE | mean | ±SE | mean | ±SE |  |  |
| *Control* |  |  |  |  |  |  |  |  |  |  |
| **α-copaene** | 0.59 | 0.13 | 0.67 | 0.17 | 0.51 | 0.09 | 1.21 | 0.22 | 6.11 | 0.11 |
| **E-β-caryophyllene** | 0.44 | 0.06 | 0.35 | 0.06 | 0.32 | 0.05 | 0.78 | 0.15 | 6.31 | 0.10 |
| **β -sesquiphellandrene<sup>N</sup>** | 1.36 | 0.26 | 1.16 | 0.34 | 1.01 | 0.19 | 2.96 | 0.69 | 5.53 | 0.14 |
| *Induced* |  |  |  |  |  |  |  |  |  |  |
| **DMNT** | 3.74 | 0.79 | 3.44 | 0.80 | 3.51 | 1.46 | 4.27 | 0.94 | 0.98 | 0.80 |
| **Indole** | 7.79 | 2.55 | 5.29 | 1.40 | 7.64 | 4.00 | 6.85 | 1.68 | 1.13 | 0.77 |
| **α-copaene** | 0.75ab | 0.16 | 0.60a | 0.11 | 1.19bc | 0.25 | 1.35c | 0.12 | 9.10 | **0.03** |
| **E-β-caryophyllene** | 0.45a | 0.08 | 0.40a | 0.07 | 0.79b | 0.18 | 0.92b | 0.13 | 9.00 | **0.03** |
| **(E)-β-bergamotene** | 1.33 | 0.34 | 1.01 | 0.20 | 1.54 | 0.49 | 1.37 | 0.22 | 1.97 | 0.58 |
| **E-β-farnesene** | 4.97 | 1.45 | 2.81 | 0.85 | 5.37 | 1.90 | 4.32 | 1.03 | 1.48 | 0.69 |
| **β -sesquiphellandrene<sup>N</sup>** | 2.08b | 0.44 | 1.57ab | 0.32 | 0.80a | 0.14 | 3.68c | 0.45 | 13.90 | **<0.001** |

### Effects of plant variety, nitrogen availability and herbivory on phytohormone and hydroxycinnamic acid accumulation

To further investigate the effect of genotype, nitrogen and herbivore attack on plant defenses, the concentrations of the phytohormones JA, OPDA, JA-Ile, SA, ABA and IAA were measured together with the hydroxycinnamic acids caffeic, ferulic and chlorogenic acid. The concentration of JA-Ile, JA, and SA, was significantly influenced by herbivory and time point (Fig. 3, models in Supplementary material Table S.1). Interestingly, feeding by the herbivore *Z. scutellaris* significantly repressed JA-Ile and JA, as mean levels of JA-Ile and JA in herbivore-damaged plants was lower than in their respective undamaged controls (Fig 3). This trend was also significant but not as clear for SA and ABA accumulation after herbivory by maize leafhoppers (Fig. 3, models in Supplementary material Table S.1). Hormone concentrations were similar among genotype per nitrogen treatments at all time points with the exception of (i) SA levels that were lower in Carolight^R^ relatively to M37W (Fig 3) and (ii) OPDA accumulated in higher concentrations in plants when grown under high nitrogen (Fig 3).

**Fig 1.**
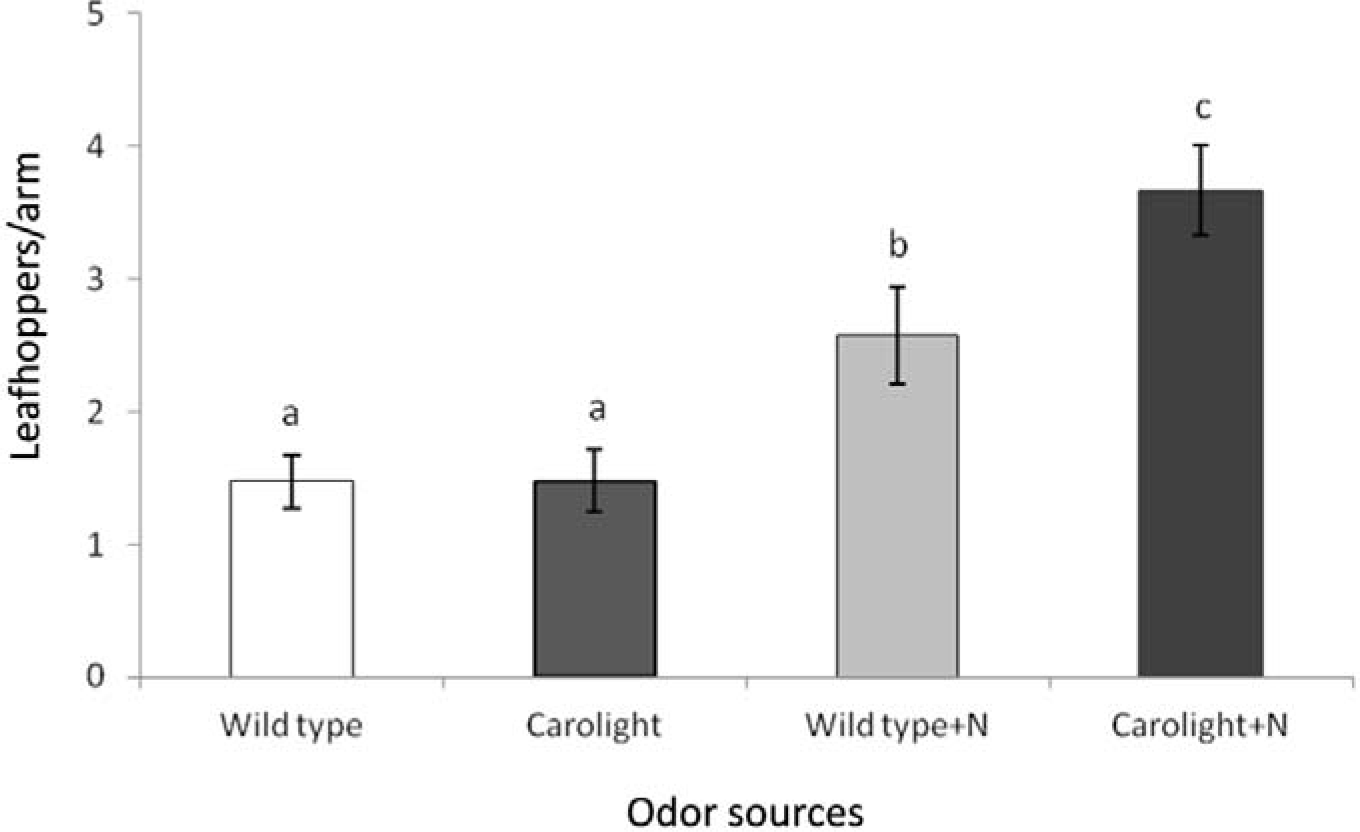
Choice of maize volatiles by leafhopper *Z. scutellaris* on the olfactometer. Tested plants consisted of Wild type and Carolight^R^ plants grown under control or surplus nitrogen (+N) conditions. Different letters indicate differences between treatments (α=0.05).

**Fig 3.**
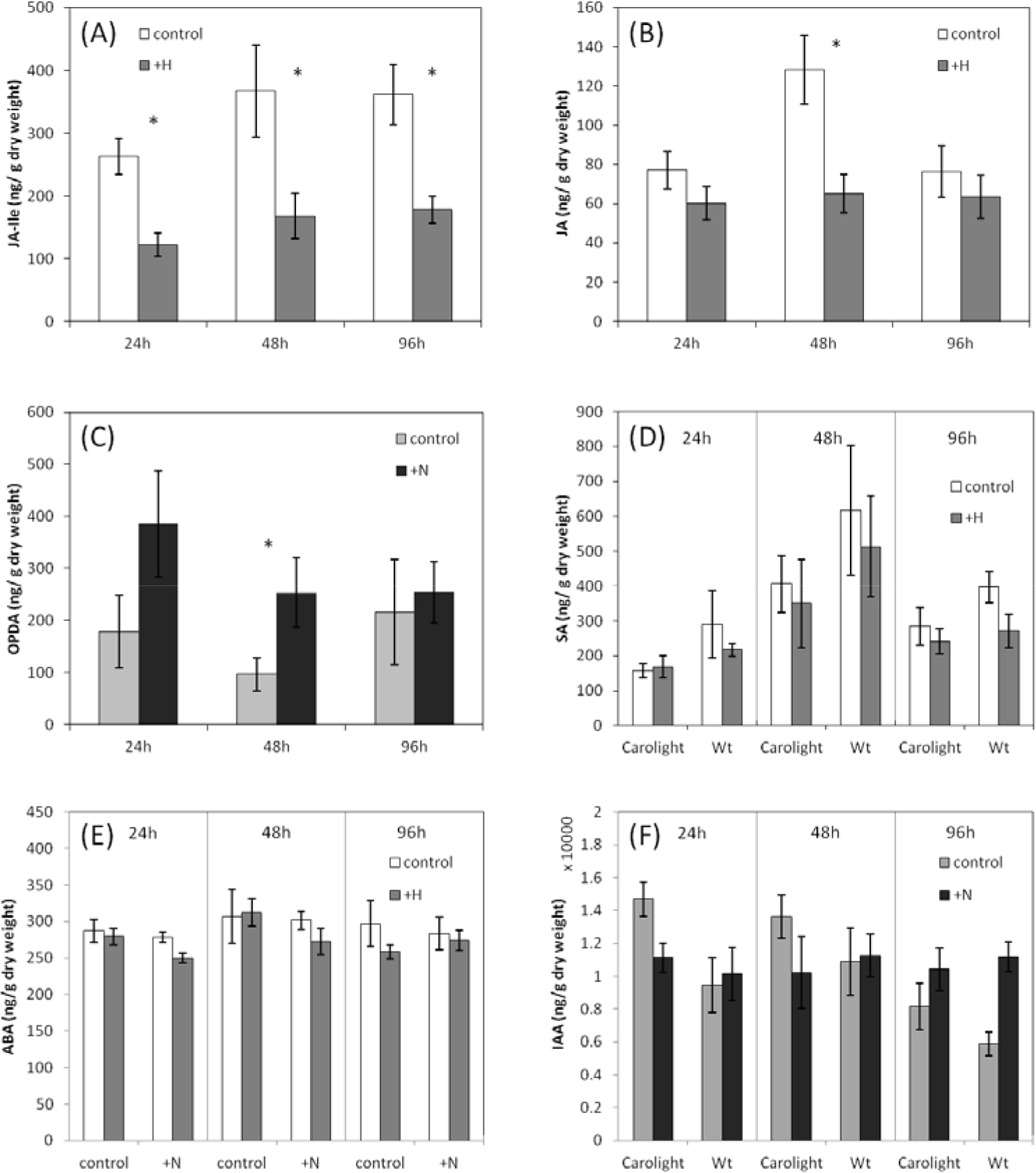
Hormonal content (ng/g dry weight) in Carolight^R^ and Wild type plants grown under two different nitrogen regimes (control and +N) upon *Z. scutellaris* feeding. Control and herbivore-damaged plants (+H) were collected at different time points (24h, 48h and 96h after herbivore feeding), and phytohormone levels were determined in freeze-dried material by UPLC-MS. The experiment was replicated 3 times with similar results. Full factorial models combining data of the three experiments are available in Supplementary material Table S.1. The results shown are mean (±SE) hormone levels of one experiment. Asterisks indicate differences among treatments (non-parametric Kruskal-Wallis test).

Overall, caffeic and chlorogenic acid concentrations were up to 2-fold lower in Carolight^R^ than in the wild type (Fig 4). Caffeic acid concentration also depended on herbivory, time point and time point per nitrogen interaction (Fig. 4 Supplementary material Table S.1), whereas chlorogenic acid accumulation varied greatly between nitrogen regimes with its concentration practically doubling under control versus high nitrogen treatments (Fig 4). No consistent differences were detected for ferulic acid accumulation for any of the factors (Supplementary material, Table S.1).

**Fig 4.**
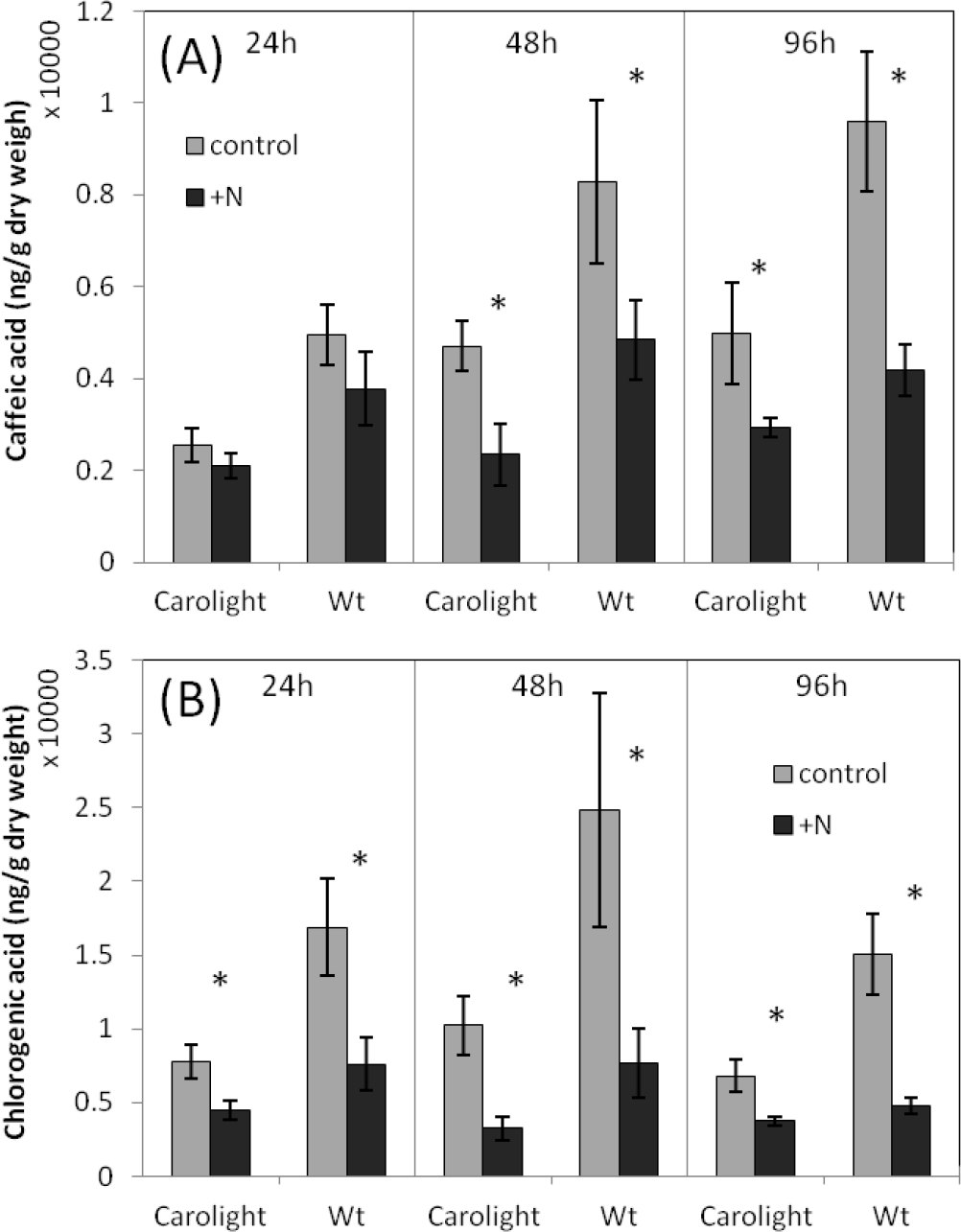
Caffeic acid (A) and chlorogenic acid (B) content (ng/g dry weight) of Carolight^R^ and Wild type plants grown under two different nitrogen regimes (control and +N). Plants were collected at different time-points (24h, 48h and 96h after the start of the experiment), and caffeic and chlorogenic acid levels were determined in freeze-dried material by UPLC-MS. The experiment was replicated 3 times with similar results. Full factorial models combining data of the three experiments are available in Supplementary material Table S.1. The results shown are mean (±SE) hormone levels of one experiment. Asterisks indicate differences among treatments (non-parametric Kruskal-Wallis test).

### Effects of plant variety, nitrogen availability and herbivory on the metabolite fingerprint

In total 4271 and 2002 markers were detected in ESI (+) and ESI (-) mode, respectively. Overall, nitrogen availability was the main factor contributing to the observed chemotypes (perMANOVA, R^2^=0.124, p=0.001), followed by genotype (R^2^=0.038, p=0.030) and herbivory (R^2^=0.034, p=0.048) while interactions of the experimental factors were non-significant. An unsupervised approach (PCA) showed that nitrogen metabolites from plants subjected to control and high nitrogen treatments clearly grouped in the first two PCs (Fig. 5), independently of the plant genotype and herbivore treatment. In contrast, genotype and herbivory related profiles could not be separated by PCA. However, a supervised partial least squares discriminant analysis (PLS-DA) model separated (i) nitrogen regimes (ii) maize genotypes, and (iii) healthy and herbivore damaged plants (Table 3, validated through CV-ANOVA). These PLS-DA models were used to identify the metabolites showing the maximum difference between treatments with VIP values >2 (Table 3), and subsequently the selected metabolites for each experimental factor (variety, nitrogen and herbivory) were screened for putative identification using the pathway tool from MarVis 2.0 software (Kaever et al 2014) (Table 3, Table 4). Mean intensities of the markers plant genotype, nitrogen and herbivory by *Z. scutellaris* are represented in Supplementary material (Fig. S6, S.7 and S.8).

**Fig 5.**
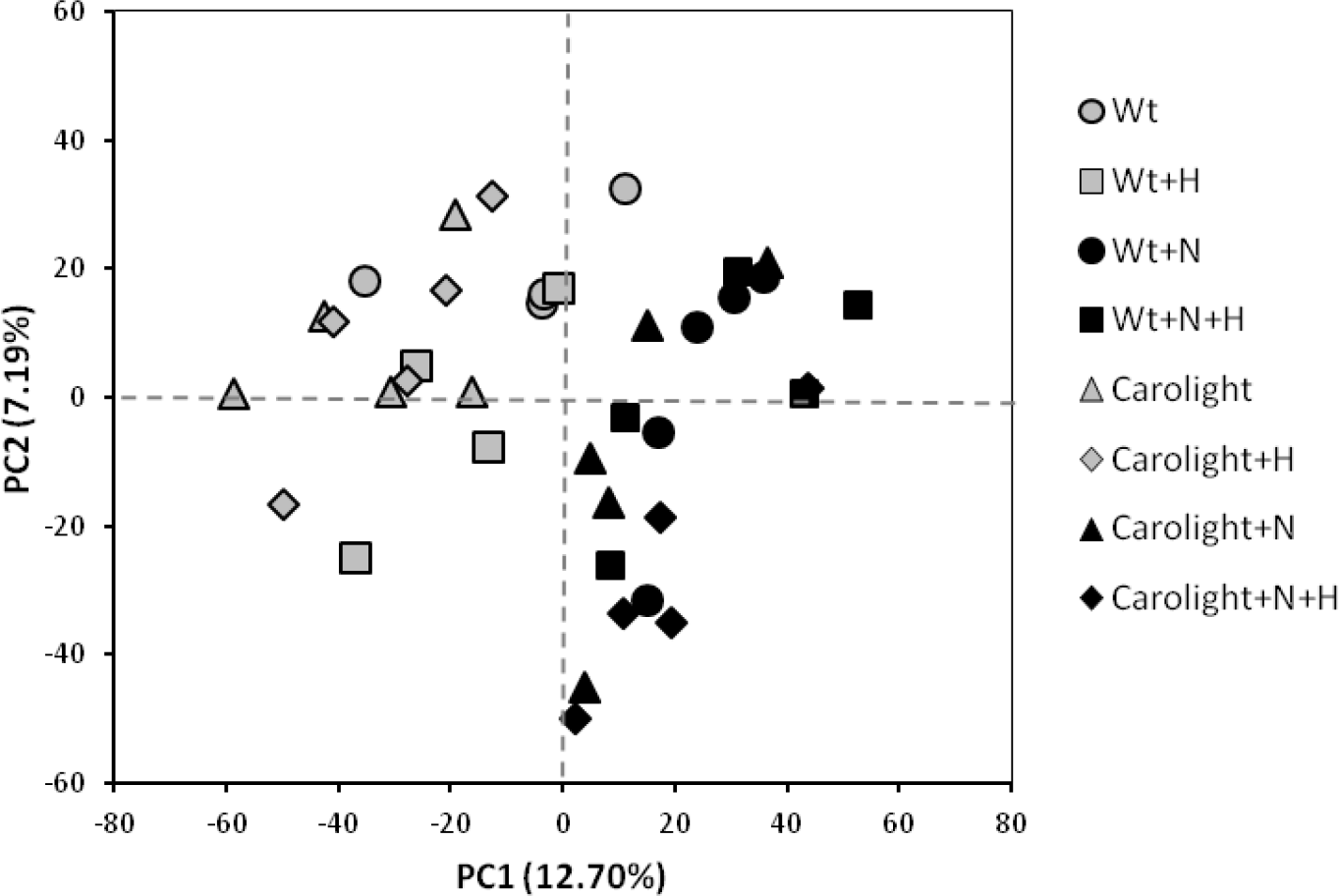
Principal component analysis (PCA) showing groups generated from signals obtained in ESI+ and ESI− by non-targeted analysis. Carolight^R^ and Wild type (Wt) plants were grown under two different nitrogen regimes (control and +N) upon *Z. scutellaris* feeding (control and +H).

**Table 3.** PLS-DA models of the factors nitrogen, genotype and herbivory. For each factor, the number of ions with a VIP>2 for the best model and the number of Marvis Pathway IDs are specified.

| Factor | Components | R<sup>2</sup> | Q<sup>2</sup> | CV-ANOVA |  | VIP>2 | MarVis ID |
| --- | --- | --- | --- | --- | --- | --- | --- |
| Nitrogen | **1** | **0.875** | **0.744** | **F<sub>2,35</sub>=50.89** | **p=4.38x10<sup>-11</sup>** | **405** | **48** |
|  | 1+1 | 0.968 | 0.999 | F<sub>4,33</sub>=34.80 | p=2.08x10<sup>-11</sup> |  |  |
| Genotype | **1+1** | **0.956** | **0.581** | **F<sub>4,33</sub>=7.87** | **p=0.00014** | **69** | **19** |
|  | 1+1+1 | 0.993 | 0.803 | F<sub>6,31</sub>=6.25 | p=0.00022 |  |  |
| Herbivory | **1+1** | **0.967** | **0.486** | **F<sub>4,33</sub>=3.97** | **p=0.0097** | **163** | **15** |
|  | 1+1+1 | 0.99 | 0.634 | F<sub>6,31</sub>=2.79 | p=0.027 |  |  |

**Table 4.** Metabolites with higher loadings on the best PLS-DA models (VIP>2) for the factors Nitrogen, Genotype and Herbivory that could be tentatively identified. Metabolites in A were identified by comparison of the MS/MS to online databases, while metabolites in B were assigned after comparing the accurate mass to reference compound databases. Mean abundances of the matabolites can be found in Supplementary material Fig. S.6, S.7 and S.8. Pathways were assigned using the pathway tool from MarVis 2.0 (Kaever et al., 2014).

| A. Identified | Mass neutral | ESI | RT (min) | Factor | Pathway |
| --- | --- | --- | --- | --- | --- |
| Kynurenic acid | 189.0431 | + | 0.99 | Herbivory (H↑) | Tryptophan-kynurenine pathway |
| 11-trans-LTD4 | 496.2592 | - | 3.43 | Herbivory (H↑) | Leukotriene- Arachidonic acid metabolism |
| 19-HETE/ 20-HETE | 320.236 | + | 2.06 | Genotype (Wt↑) | Arachidonic acid metabolism |
| Thiamine | 265.1153 | + | 0.86 | Genotype (Wt↑) | Vitamin and cofactor - Thiamine metabolism (primary metabolism) |
| Phytosphingosine | 317.2932 | + | 4.41 | Genotype (Wt↑) | Sphingolipid metabolism (primary metabolism) |
| L-Tryptophan | 204.0897 | - | 0.93 | Nitrogen (+N↑) | Tryptophan pathway (primary metabolism) |
| DIMBOA-Glu | 373.1009 | - | 1.46 | Nitrogen (+N↑) | Benzoxazinoid biosynthesis |
| B. Assigned to | Mass neutral | ESI | RT (min) | Factor | Pathway |
| Sinapoyl malate | 340.0791 | - | 2.73 | Herbivory (H↓) | Biosynthesis of phenylpropanoids |
| Porphobilinogen | 226.0965 | - | 0.76 | Herbivory (H↑) | Porphyrin and chlorophyll metabolism |
| (-)-Jasmonoyl-L-isoleucine | 323.2097 | + | 4.97 | Herbivory (H↓) | Biosynthesis of plant hormones |
| A-tocopherol | 430.3777 | + | 5.92 | Herbivory (H↑) | Biosynthesis of plant secondary metabolites |
| 2,3-Dihydroxybenzoate | 154.0263 | - | 1.44 | Herbivory (H↑) | Biosynthesis of phenylpropanoids |
| Coniferol | 180.0788 | + | 0.84 | Genotype (Wt↑) | Biosynthesis of phenylpropanoids |
| Cis-hinokiresinol | 252.1195 | - | 0.84 | Genotype (Wt↑) | Biosynthesis of phenylpropanoids |
| Unknown flavonoid | 306.0775 | - | 1.36 | Genotype (Wt↓) | Flavonoid biosynthesis |
| Unknown flavonoid | 578.1634 | + | 1.78 | Genotype (Wt↓)<br>Nitrogen (+N↓) | Flavone and flavonol biosynthesis |
| Unknown flavonoid | 610.1544 | + | 1.79 | Genotype (Wt↓)<br>Nitrogen (+N↓) | Flavone and flavonol biosynthesis |
| Unknown flavonoid | 448.1016 | +/- | 1.59/2.05 | Nitrogen (+N↓) | Flavone and flavonol biosynthesis |
| Unknown flavonoid | 594.1579 | - | 1.61 | Nitrogen (+N↓) | Flavone and flavonol biosynthesis |
| Unknown flavonoid | 286.0483 | + | 1.99 | Nitrogen (+N↓) | Flavone and flavonol biosynthesis |
| Unknown flavonoid | 464.0949 | - | 1.86 | Nitrogen (+N↓) | Flavone and flavonol biosynthesis |
| Unknown flavonoid | 592.1802 | + | 2.73 | Nitrogen (+N↓) | Flavone and flavonol biosynthesis |
| Zeatin/Pantothenate | 219.1113 | + | 0.8 | Nitrogen (+N↓) | Biosynthesis of plant hormones |
| HBOA | 149.0471 | - | 0.73 | Nitrogen (+N↑) | Benzoxazinoid biosynthesis |
| DHBOA/ DIBOA-Glc | 343.0891 | - | 1.27 | Nitrogen (+N↑) | Benzoxazinoid biosynthesis |
| Dhurrin | 311.1002 | - | 0.73 | Nitrogen (+N↑) | Cyanoamino acid metabolism |

## DISCUSSION

We addressed the hypothesis that a nutritionally enhanced maize (Carolight^R^), similar in terms of biomass and yield to its wild type line M37W (Zanga *et al.*, 2016), will also be equivalent in terms of plant-insect interactions. Evaluating Carolight^R^ and wild type genotypes under contrasting nitrogen levels allowed for (i) a broader characterization of the resulting chemotypes and their impact on insect behavior/performance; and (ii) a comparative analysis of the impact of experimental factors (nitrogen, genotype, herbivory) on the final chemotypes. We demonstrated that nitrogen availability is the main factor determining herbivore preference and the metabolite fingerprint in Carolight^R^ and M37W, followed by the introduced traits and herbivory. There were no significant effects of nitrogen x genotype or herbivory x genotype interactions, suggesting that both genotypes behaved similarly when grown under the same nitrogen conditions.

### Insect abundance and performance on Carolight^R^ in contrasting nitrogen availability conditions

Overall, the community of herbivores was similar for both Carolight^R^ and M37W genotypes. Yet in the case of Hemiptera (leafhoppers, planthoppers and aphids) higher abundances were detected in plots with high nitrogen while only the leafhopper *Z. scutellaris* nymph abundances were significantly higher for M37W. Nitrogen is one of the most frequently used fertilizers in agricultural production and is known to exert a variety of bottom-up effects and potentially alter tritrophic interactions through various mechanisms (Chen *et al.*, 2010), especially for herbivorous Hemiptera (Butler *et al.*, 2012). Hemipterans are insects with a high potential sensitivity to plant quality as they have been reported to prefer and perform better on some genotypes or on plants that differ in quality in terms of nutritional requirements (e.g. nitrogen content), physical or chemical plant defense (e.g. Kallenbach et al., 2011; Zytynska and Preziosi, 2011). A number of reports suggest that herbivore Hemiptera (especially leafhoppers and aphids) are more abundant and/or perform better on Bt maize lines compared to their corresponding near isogenic counterparts (Lumbierres *et al.*, 2004, 2010; Pons *et al.*, 2005; Obrist *et al.*, 2006; Virla *et al.*, 2010; Rauschen *et al.*, 2011). The underlying mechanism(s) responsible for such differences have not been attributed to specific factors, rather to pleiotropic effects. Pleiotropic effects reported for Bt maize that might influence Hemipteran densities are higher lignin content in the stem of Bt plants (Saxena and Totzky, 2001), reduced amount of VOC emission in a Bt line (Turlings *et al.*, 2005) and sap amino acid content (Faria *et al.*, 2007).

Insect herbivores are limited by low nitrogen concentrations in food plants, and therefore herbivore performance is generally thought to be positively related to increases in nitrogen content in plants (Awmack and Leather, 2002; Behmer, 2009; Butler *et al.*, 2012). The performance of *Z. scutellaris* nymphs was similar when fed on Carolight^R^ and M37W grown under control and high nitrogen levels. This result was unexpected as we hypothesized that nitrogen availability would be the main factor contributing to adult final weight as a proxy for reproductive fitness. However, female leafhoppers preferred maize plants grown under high nitrogen in the olfactometer test, and even preferred Carolight^R^ over M37W when plants were grown under high nitrogen. This fact - together with field data on leafhopper abundance - supports the notion that host plant quality (resulting from enhanced nitrogen fertilization) might indeed offer other advantages to the species, such as reproductive success, that are not reflected by adult body weight or duration of nymphal development. Prestidge (1982) reported an increasing oviposition of *Z. scutellaris* as the nitrogen fertilization increased in the grass *Holcus lanatus.* Therefore the lack of differences in adult weight and developmental time for maize leafhoppers in our experiments could be a product of a mismatch between adult size and fecundity in *Z. scutellaris* as it has been previously described for grasshoppers (Joern and Behmer, 1998). Several features including field abundance, plant preference and fecundity could provide the best measures of performance for *Z. scutellaris* in general.

### Maize defense responses to a mesophyll-feeding leafhopper

Plant damage together with salivary secretions of phytophagous arthropods are known to trigger plant inducible defense responses (Alborn et al. 1997, Musser et al. 2002). In turn inducible plant defenses can be major determinants of ecological interactions, and in particular defenses depending on JA and SA pathways appear to play important roles in determining community composition (Thaler *et al.*, 2001; Thaler, 2002; Kallenbach *et al.*, 2011). Hence it was vital to determine whether Carolight^R^ and its wild type parent (M37W) behave similarly in terms of constitutive profiles of JA and SA and in hormonal response when facing herbivory. Cell content feeders, such as the spider mite *Tetranychus urticae* Koch (Acari: Prostigmata) and the thrips *Frankliniella occidentalis* (Pergande) (Thysanoptera:Thripidae) usually stimulate JA-inducible genes upon attack (Vos *et al.*, 2005), although there are reports that confirm the activation of both SA- and JA-inducible genes (Kant *et al.*, 2004; Kawazu *et al.*, 2012). Given that typhlocybine leafhoppers such as *Z. scutellaris* feed on the mesophyll using a sawing laceration strategy (Marion-Poll et al. 1987, Backus et al. 2005) we predicted that feeding by the leafhopper would activate either the JA and/or SA pathways. Interestingly feeding by this herbivore appears to decrease the constitutive levels of JA and SA on maize plants, and this was reflected in that phytohormone levels in leafhopper damaged plants were similar or even lower than the constitutive levels in healthy control plants, in particular those of JA-Ile.

Suppression of plant defenses is a well-known phenomenon in plant pathogens such as pathogenic bacteria, rust fungi, oomycetes, viruses, and herbivores such as nematodes and spider mites (reviewed by Kant et al. 2015 and Zhang et al. 2017). Spider mite *Tetranychus evansi* suppresses both JA and SA dependent defenses in tomato enhancing their performance (Sarmento *et al.*, 2011; Alba *et al.*, 2015). In the case of insects the majority of cases of plant defense suppression has been attributed to JA-SA hormonal crosstalk (Walling, 2000; Zhang *et al.*, 2017) and not to a direct blocking of JA or SA defenses. However, recently aphids and mites have been reported to deliver effectors when feeding as a strategy to overcome host-plant defenses and improve their fitness (Hogenhout and Bos, 2011; Kant *et al.*, 2015; Mugford *et al.*, 2016; Villarroel *et al.*, 2016). In our experimental system, *Z. scutellaris* - by feeding and oviposition - suppresses JA and does not induce SA in maize plants, and hence hormonal suppression appears to occur independently of SA-JA cross talk (as was the case for *T. evansi*). Non-targeted metabolomics fingerprinting allowed the identification of markers of herbivory by *Z. scutellaris*, which opens a door to further research on the potential effectors delivered by the leafhopper and the mechanism of defense suppression.

Defense manipulation by maize leafhopper impaired phytohormone accumulation in the plant without disturbing plant indirect defense by means of herbivore induced plant volatile emission. Previous work showed that maize plants damaged by ten *Z. scutellaris* adults emitted a similar amount of volatiles than plants damaged by the five 2nd instar *Spodoptera littoralis*, and that the predatory anthocorid *Orius majusculus* was innately attracted towards the volatile blend (Ardanuy *et al.*, 2016). We hypothesize that the suppression of JA defenses ultimately benefits leafhopper reproduction and nymphal performance, but natural enemies will still protect the plant through top-down control. However, defense manipulation by maize leafhoppers might also have consequences for subsequent colonizing herbivores since maize plants with suppressed defenses might promote the performance of co-occurring herbivores (Stam *et al.*, 2014; Kant *et al.*, 2015).

### Nitrogen determines the chemical defense attributes of Carolight and M37W

Evaluating Carolight^R^ and wild type genotypes in contrasting nitrogen conditions allowed for a comparative analysis of the impact of experimental factors (nitrogen, genotype, herbivory) on the final maize chemotypes. We demonstrated that nitrogen is the main factor determining the metabolite fingerprint in Carolight^R^ and M37W, followed by the introduced traits and herbivory. Our results corroborate the work of Coll et al. (2010) where transcript analysis in two maize Bt(Cry1Ab trait)/wild type pairs in the field indicated that differences between lines (genetic background) exerted the highest impact on gene expression patterns, followed by nitrogen availability, while the Cry1Ab trait had the lowest impact. Barros et al. (2010) compared two GM maize pairs - Bt (Cry1Ab) and glyphosate tolerant - using transcriptome, proteome, and metabolome profiling and reported that the environment affected gene expression, protein distribution, and metabolite content more strongly than the introduced traits. Our results are therefore consistent with the literature and show that environmental factors (e.g. field location, sampling time during the season or at different seasons, mineral nutrition) consistently exert a greater influence on crop lines than the genetic modification itself (reviewed by Ricroch et al. 2011).

In general, nitrogen fertilization increases plant growth and reproduction, decreases concentrations of carbon-based secondary compounds (e.g. phenolics and terpenoids), and increases nitrogenous compounds (Koricheva *et al.*, 1998; Lou and Baldwin, 2004; Scheible *et al.*, 2004; Hermans *et al.*, 2006; Kusano *et al.*, 2011). Nitrogen levels influenced Carolight^R^ and M37W phenotypes at the metabolite level substantially, including compounds involved in direct and indirect plant defenses. Of the potential 405 markers with a VIP>2 only few were putatively identified. Some of these are secondary metabolites and contribute to the plant’s constitutive defense as flavonoids or hydroxamic acids (benzoxazinoids) (Table 4, Supplementary material Fig. S.6). Targeted analysis of defense metabolites showed that chlorogenic acid greatly varied with nitrogen treatments - at higher concentrations in plant tissues when nitrogen was limiting - but also with the plant genotype - M37W had higher levels of both chlorogenic and caffeic acids. Higher concentration of constitutive phenolics in plants under low nitrogen is consistent with results in *Nicotiana atenuata* (Lou and Baldwin, 2004) and tomato (Stout *et al.*, 1998). Carolight^R^ accumulated up to 2-fold lower amounts of plant hydroxycinnamic acids (caffeic and chlorogenic acids) depending on the nitrogen treatment and time-point, and higher amounts of lignans (especially at low nitrogen) than the wild type. This suggests an effect of the genotype on the phenylpropanoid biosynthetic pathway. In addition, phenolics in the form of unidentified flavonoids were more abundant in control nitrogen maize plants (Table 4, Supplementary material Fig. S.6).

Metabolite fingerprinting showed that nitrogen surplus increased the accumulation of tryptophan in plants, which we identified as a marker of high nitrogen treatment. Tryptophan serves as precursor of a broad variety of nitrogen-containing aromatic secondary metabolites, such as hydroxamic acids (Fig. S.6), which play crucial roles in plant defense against herbivore feeding (Niemeyer, 2009; Balmer *et al.*, 2013). Higher levels of constitutive phenolics and hydroxamic acids would theoretically increase plant tolerance towards herbivores, as increased levels of these secondary compounds have been associated to reduced herbivory (Mithöfer and Boland, 2012; Balmer *et al.*, 2013). Olfactometer plant choice might indicate a preference towards plants with lower concentration of phenolics and higher concentration of hydroxamic acids in the plant; however, it fails to explain higher abundance of maize leafhopper nymphs in wild type plots in the field.

The VOC blend was also modified by nitrogen availability: a higher concentration of the sesquiterpenes α-copaene and E-β-caryophyllene was detected for Carolight^R^ and M37W plants under higher nitrogen. This might explain leafhopper preference towards plants grown under higher nitrogen levels. These results contrast with previous findings (Schmelz et al. 2003) which reported higher VOC emission in maize with limited nitrogen availability, though differences could be explained by maize varieties or by the source of nitrogen used in each study. While we applied nitrogen as both nitrate and ammonium, in the later study nitrogen was applied as nitrate. Nitrogen deficient soybean plants emitted the same range of herbivore-induced VOCs as control plants, but quantitative changes occurred in the release of the main compound β-farnesene and two other volatiles ((Z)-3-hexenyl-α-methylbutyrate and β-bergamotene) (Winter and Rostás, 2010) and no differences were detected in *Nicotiana attenuata* (Lou and Baldwin, 2004).

Carolight^R^ damaged plants grown under high nitrogen emitted a larger amount of β-sesquiphellandrene; however the change in the volatile blend did not influence the community of natural enemies in the field. A blend of VOCs that varies in the composition or quantity of its components may constitute a signal with altered information content and may potentially modify the host finding behavior of herbivores and natural enemies, as it is the case for the maize leafhopper *Z. scutellaris*, which prefers Carolight^R^ to the wild type when grown under high nitrogen. It remains unclear whether leafhoppers respond in a dose-dependent manner to the total blend of VOCs or if other compounds at doses too small to detect (D’Alessandro et al. 2006) triggered leafhopper preference in the olfactometer.

### Conclusion

We show the separate and interactive effects of nitrogen availability and genotype on the arthropod community and on the performance and behavior of a herbivore, and correlated these changes to constitutive and inducible maize defenses. We conclude that: (i) nitrogen availability greatly shapes maize metabolism, and the resulting plant chemotypes, and promotes *Z. scutellaris* preferences through the emission of a more attractive blend of VOCs; (ii) feeding by *Z. scutellaris* suppresses the accumulation of JA-Ile, JA and SA, while triggering the emission of herbivore-induced plant volatiles; and (iii) that the minor differences detected among Carolight^R^ and its wild-type counterpart in the phenylpropanoid pathway do not substantially alter aboveground plant-arthropod interactions.

## ACKNOWLEDGMENTS

We would like to thank Michael Gaillard and Daniel Maag for help in sample preparation, Gemma Camañes for advice on nitrogen treatments, Diana LaForgia for help in arthropod sampling, Joan Safont, Carmen López and Matilde Eizaguirre for providing *Z. scutellaris*, Daniela Zanga for discussions on Carolight^R^ agronomic performance, Victor Flors and the Metabolic Integration and Cell Signaling Group for discussion on results and data analysis, and all the members of the FARCE and E-Vol groups in Neuchâtel for their collaboration. A.A. was funded with an FPU scholarship and a visiting grant to FARCE laboratory from the Ministerio de Educación, Spain. The work was partially funded by the Spanish Government projects AGL2011-23996 and AGL2014-53970-C2-1-R, Agrotecnio (Fundació Centre de Recerca en Agrotecnologia, VitaMaize 01 project) and Fundació la Caixa (Grant P13005).

